# New Interactive Machine Learning Tool for Marine Image Analysis

**DOI:** 10.1101/2023.10.12.562046

**Authors:** H. Poppy Clark, Abraham George Smith, Daniel Mckay Fletcher, Ann I. Larsson, Marcel Jaspars, Laurence H. De Clippele

**Affiliations:** Marine Biodiscovery Centre, Department of Chemistry, University of Aberdeen, Old Aberdeen, AB24 3DT, Scotland, United Kingdom; Department of Computer Science, University of Copenhagen, Universitetsparken 1, 2100 Copenhagen, Denmark; Rural Economy, Environment and Society, Scotland’s Rural College, West Mains Road, Edinburgh, EH9 3JG, Scotland, United Kingdom; Tjärnö Marine Laboratory, Department of Marine Sciences, University of Gothenburg, Strömstad, Sweden; School of Biodiversity, One Health & Veterinary Medicine, University of Glasgow, Bearsden Road, G61 1QH, Scotland, United Kingdom

## Abstract

Due to advances in imaging technologies, the rate of marine video and image data collection is drastically increasing. Often these datasets are not analysed to their full potential as extracting information for multiple species, such as their presence and surface area, is incredibly time-consuming. This study demonstrates the potential of a new open-source interactive machine learning tool, RootPainter, to analyse large marine image datasets quickly and accurately. The tool was initially developed to measure plant roots, but here was tested on its ability to measure the presence and surface area of the cold-water coral reef associate sponge species, *Mycale lingua*, in two types of underwater image data: 18,346 time-lapse images and 1,420 remotely operated vehicle video frames. New corrective annotation metrics integrated with RootPainter, such as dice score and species area error, allow for the objective assessment of when to stop model training and reduce the need for manual model validation. Three highly accurate *Mycale lingua* models were created using RootPainter, as indicated by their average dice score of 0.94 ± 0.06. Model transfer and optimisation aided in the production of two of these models, increasing analysis efficiency from 6 to 16 times faster than manual annotation in Photoshop, for underwater observatory images. Sponge and surface area measurements were extracted from both datasets allowing future investigation of sponge behaviours and distributions. This study demonstrates that interactive machine learning tools and model sharing have the potential to dramatically increase image analysis speeds, collaborative research, and our collective knowledge on spatiotemporal patterns in biodiversity.

## INTRODUCTION

Large image datasets enable detailed and long-term studies of underwater species, providing a vital tool for the ecological investigation of deep-water species [1–3]. However, extracting information of interest from image datasets, such as species presence or size, can be prohibitively time-consuming [4–6]. This problem is exacerbated for more complex images, such as those captured by mobile cameras on remotely operated vehicles (ROVs), or automated underwater vehicles (AUVs), where lighting and focus may vary compared to stationary underwater cameras or fixed observatories [7,8]. There has therefore been a trend to develop bespoke machine learning algorithms to extract information from a given image dataset; the most successful of these result from collaborations between marine and computer scientists [4,9,10]. As machine learning algorithms are non-trivial to construct and apply, their accessibility to individuals without experience in scientific programming languages is limited, creating a barrier to model sharing that is increasing the redundancy of work being completed by image analysis groups.

Pre-developed, user-friendly, and widely applicable machine learning tools may present a solution to some of these issues. They allow individuals with no machine learning or coding skills to develop models through training a pre-existing and adaptable ‘base’ neural network. The process to train models can vary depending on the tool employed, but their complexity is often masked through user interfaces. Improved accessibility of machine learning for image analysis increases the rate and range of measurements that can be extracted from image data, facilitating a variety of ecological investigations, as has previously been demonstrated through manual analyses. Extracting the length, perimeter, or area of species from images allows estimation of their size, and/or growth rates [11–15] and extracting count data can be used in species abundance [16], or biodiversity estimates [17]. Previously unknown species behaviour traits can be revealed by tracking individuals through extraction of their x,y coordinates [2], and extraction of ‘global shape measures’ such as the eccentricity, or curvature, of subjects of interest may allow investigation of morphological diversity [18]. Combining analyses can increase the value of information obtained further, for example extracting both species abundance and individual areas allows estimation of biomass [15,19,20], and investigating sessile species size alongside local biotic or abiotic factors may inform on their behaviour [21,22].

These measurements have therefore been key targets in marine machine learning studies. Whilst algorithms capable of automated species detection have been developed [4,23–25], they are yet to be utilised to produce biodiversity estimates. This may be the result of inherent difficulties associated with automated species detection, such as the need for each species to be annotated a sufficient number of times in the training data to be detected when the model is subsequently applied [6]. Conversely, development of machine learning algorithms capable of predicting the area of sessile organisms from marine images has led to successful investigation of behaviour such as cold-water coral feeding [26,27], and sponge contractions [9,28,29]. Some algorithms have taken a step further, attempting to simultaneously investigate the areas of multiple species to study community structure and ecosystem functions [10]. However, as Purser *et al.* found, a major barrier to this is the differing visual complexity of species; their model performed well when estimating live cold-water coral density but struggled to accurately assess sponge coverage. Future investigations were thus completed through manual annotations [30]. Given these challenges, an approach where simultaneous extraction of multiple measurements for one species from image data may provide an alternative solution to increase the speed and complexity of ecological conclusions possible in these studies. There is therefore a significant need for an accessible and multi-functional machine learning tool to analyse marine image data.

RootPainter is an open-source and user-friendly graphical user interface-based software tool enabling the rapid training of convolutional neural networks via corrective annotation [31]. It is classified as an interactive machine learning tool as the user is part of the training feed-back loop involved in model development. RootPainter was developed to investigate root length and the presence of soil voids (bipores) in soil images, with the production of a successful model being achievable within one working day. Users are not required to possess a high-powered Graphics Processing Unit (GPU), or any coding competencies and models can be transferred between projects and users. Internally, RootPainter uses a variant of the general purpose U-Net convolutional neural network [32], that has been found to be effective for many tasks. As RootPainter introduces no requirements on the type of structures that a model can be trained to detect, its applications are not limited to plant images. However, the innate complexity associated with marine images (due to suspended matter affecting image clarity and the uneven illumination of scenes with artificial lighting in the depths) may increase model training time compared to images from controlled (laboratory) conditions. Previous studies employing automated marine image analysis have relied heavily on image pre-processing to diminish the complexity of their image datasets [9,26]. This can involve denoising or brightness/contrast/colour normalisation and is one of the most time-consuming stages of the analysis pipeline [10,33]. Reliance on non-trivial image pre-processing not only reduces the accessibility of machine learning algorithms and the transferability of models, but also limits the processing pipeline to application on that specific dataset. In order to avoid these issues image pre-processing is not required to produce successful models with RootPainter [34–37].

This study, therefore, investigates the suitability of RootPainter to marine image analysis. The potential of RootPainter to combat key challenges in the field was explored through testing its ability to identify and predict the surface area of a known difficult target for machine learning algorithms, the deep-sea sponge *Mycale lingua* [10]. Additionally, RootPainter’s capabilities with images of varying complexity were assessed through comparison of model performance for static timelapse images from an underwater observatory, and frames extracted from ROV videos.

## MATERIALS AND METHODS

### Study Sites, Underwater Imagery and Data Availability

Data was used from two separate locations in Norway (Figure 1a). Time-lapse imagery was captured at the cabled Lofoten Vesterålen (LoVe) Ocean Observatory at 240 m depth [38], and ROV videos were recorded at the Tisler reef, between 70-160 m depth [39].

**Figure 1:**
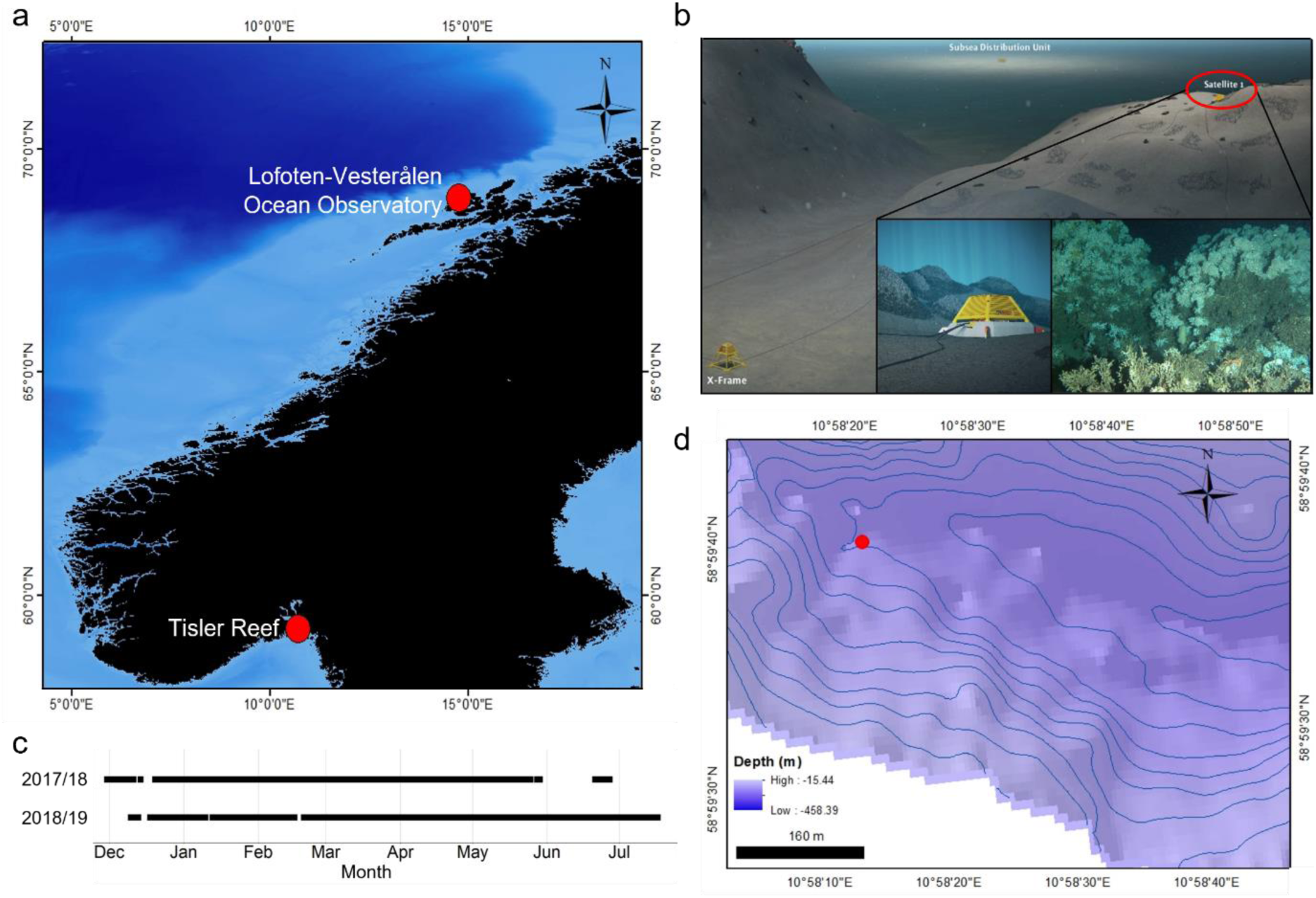
(a) Map of Norway highlighting the locations of the LoVe Ocean Observatory and Tisler reef, (b) Sub-sea layout of the LoVe Ocean Observatory including the satellite 1 structure responsible for collecting data used in this study and an example of the raw 5202 x 3464 pixel image output from satellite 1 during 2017-19 (adapted from [38]), (c) Image data availability from the LoVe observatory between 2017 and 2019, (d) Bathymetry map of the Tisler reef, with the 2021 ROV survey area pin-pointed.

The LoVe Observatory (N 68°54.474′, E 15°23.14) is in the Hola trough, a continental slope 20 km from the Lofoten Islands [38]. Sub-station satellite 1 (Figure 1b) was installed in 2017, supporting a Canon EOS 550 camera with E-TTL flash mode that captured 9,173 hourly images throughout 2017, 2018 and 2019. Data was transferred from the satellite through 450 m of subsea cable, via the X-frame, to the observatory main cable at the Subsea Distribution Unit, (Figure 1b, [38]); observatory structure maintenance resulted in data gaps during this time (Figure 1c).

The Tisler reef is found north of the Tisler Island in a 48-km-long ocean channel in the Hvaler area [7]. The research vessel Nereus, stationed at the Tjärnö Marine Laboratory was used to deploy the Ocean Modules ROV (V8 Sii, P/N: 02/00100-01, S/N: 011) to record the eastern section of the reef in 2021 (Figure 1d). A full-colour HD Hama lens camera with two Bowtech LED-K-2400 lights (2400 lumens each) was used to collect the video footage. Video signals were transmitted over an optical fibre as the ROV moved. Two laser beams, separated by 5 cm, were used as a reference to scale video frames. An Applied Acoustics Nexus Lite USBL system, running the Applied Acoustics 1329A Micro beacon provided ROV navigation data. Every 130^th^ frame was extracted from a total of 1 hour and 55 minutes of video; this minimised content overlap between frames but maximised reef coverage (ROV speed varied during the survey). A total of 1,420 images of 1920 x 1088 pixels were extracted as a result.

### Target Species

The Lofoten Vesterålen region and Tisler reef both host abundant *Desmophyllum pertusum* colonies (alternately known as *Lophelia pertusa*, Linnaeus 1758, [40]) and sponges, including *Mycale lingua* (Bowerbank, 1866). *M. lingua* was chosen as the target species to explore the capabilities of RootPainter as the complex range of colours, textures and morphologies that sponges display within a given species makes them difficult subjects for machine learning algorithms [10].

*Mycale lingua* (Bowerbank, 1866) is a Demospongiae found widely distributed across the northern hemisphere at depths of 30-2,500 m, with particularly high concentrations in the North Atlantic Ocean [41]. *M. lingua* non-selectively consumes small (<10 µm) plankton [42] and is one of the only sponge species known to successfully colonise reef areas that have high *L. pertusa* densities [10,30,43]. Other than its association with cold-water coral reefs, little is known about the behaviour of *M. lingua*; thus far there has been limited success maintaining the sponge in aquaria for extended periods of time [44]. As sponges are important components of benthic ecosystems, both in the presence and absence of *L. pertusa* reefs [30,45], understanding their distribution, biomass and behaviour could allow evaluation of factors such as their contribution to carbon-cycling in benthic environments [19].

Utilising both the LoVe Ocean Observatory and Tisler reef datasets allows exploration of the ability of RootPainter to identify *M. lingua* from a more uniform dataset (i.e. one sponge in one location) and a more diverse dataset (i.e. different *M. lingua* individuals in different locations). In this study adjoining sponge lobes were treated as one individual. *M. lingua* are known to exhibit lobed body structures [41], and without sampling it was not possible to confirm whether lobes were genetically distinct.

### RootPainter

The software program RootPainter works through three stages:

**Stage 1:** Users annotate images with clear examples until non-random model predictions are seen (requires 6-10 images).
**Stage 2:** Users switch to corrective annotation, continuing to work through the training images which have been pre-segmented (images displaying predictions) by the current model. These corrections are included in the training data, continuously improving the model until users are satisfied with its performance, which is also indicated by multiple corrective annotation metrics.
**Stage 3:** The trained model is then used to automatically process (segment) the full dataset.

The continuous feedback loop in stage 2 allows issues and anomalies to be addressed by the user that may have not been encountered in stage 1. This corrective annotation continually supplies measures of true and false, positives and negatives to the algorithm for each image. Multiple corrective annotation metrics (i.e. precision, recall, dice score and accuracy) can therefore be calculated during training without the need for separate manual annotations to validate the performance of the model. RootPainter (version 0.2.23 onwards) can also estimate the error in the area predicted by its models during training, allowing assessment of model success.

Once trained, models classify images into foreground and background, where foreground is the object of interest. These predictions are called segmentations; a visual output is provided for each image where segmentations are shown as blue highlighted regions. From these segmentations six measurements can be simultaneously extracted by RootPainter, these include count and area of regions of interest, as well as the diameter, perimeter and x,y coordinates of each discrete area. Users may also choose to extract the eccentricity (a proxy for curvature) of each discrete area. When several subjects are present within one image, separate values for their finite areas, as predicted by RootPainter, are reported.

### RootPainter Model Training

RootPainter installation and model development were completed as per the GoogleColab notebook instructions [46]. A detailed manual describing model training, specifically for marine images, is also available [47].

The LoVe Observatory images were cropped using the “magick” package in R, to form two datasets containing 2200 x 2550 and 1000 x 1964 pixel images, each containing one *Mycale lingua* individual hereafter referred to as Magnus and Mini, respectively (Figure 2). This allowed evaluation of the efficiency of RootPainter on images of different sizes, as well as the model transfer function of RootPainter within a dataset. The ROV frames were cropped to 1400 x 888 pixels using the “magick” package in R, such that the lasers were centralised, the ROV display text was removed, and the far background of each was image limited (Figure 2).

**Figure 2:**
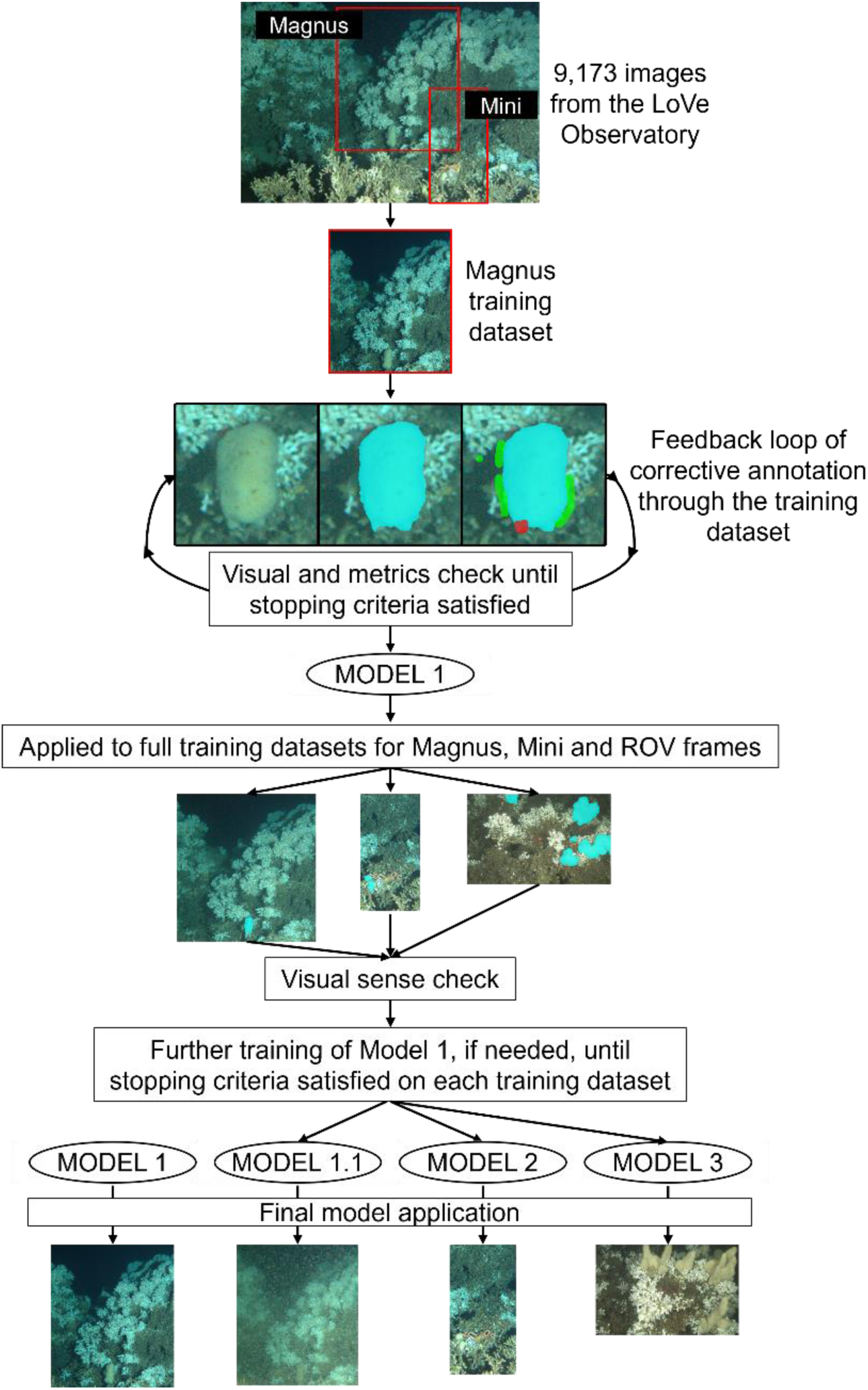
Model development workflow. Magnus and Mini from the LoVe Observatory were cropped into separate images, forming datasets of 9,173 images each. The images of Magnus from 2019 were uploaded to GoogleDrive forming a training dataset. During RootPainter model training, the algorithm presented successive random images from the training dataset, along with its prediction for that image. The user then corrected this prediction, highlighting in green pixels that should be included background and in red pixels that should be included in the foreground. The continuous visual feedback loop and accompanying metrics allowed determination of the endpoint of training, producing Model 1. This model was then applied to the Magnus, Mini and ROV frame datasets (the ROV images are shown at 1.5 times their true size, relative to the LoVe observatory, for improved visualisation). After checking the segmentation outputs, further model training was clearly required on April/May 2018/19 for Magnus, and on images from 2018 for Mini; significant further training of Model 1 was required on the ROV frames. This additional training produced Models 1.1, 2 and 3. All four models were then applied to their total respective datasets.

RootPainter was run through the free version of GoogleColab, with GoogleDrive used to sync image directories. To comply with free storage limits, one year of images from the LoVe Observatory were uploaded to GoogleDrive as the training datasets for Magnus and Mini; images of Magnus from 2019 and Mini from 2018 were used as a shift in coral rubble obscures one of the lobes of Mini in 2019. The total ROV dataset of 1,420 images was uploaded to GoogleDrive for use in training.

Once running, RootPainter presents random successive images to the user from the selected training dataset. Eight of these images were annotated with examples of foreground (species/substrate of interest, here *M. lingua*) and background (everything else in the image) before models provided non-random predictions. Subsequent segmentations were then corrected, highlighting false positives (overpredictions that should be background) in green and false negatives (underpredictions by the current model) in red (Figure 2). These corrections were incorporated into successive new and improved models.

In total five RootPainter models were produced. Model 1 was developed on images of Magnus. Additional fine-tuning of Model 1 was required on images of Magnus from April/May of 2018/19 due to a change in sponge colour/texture and turbid conditions; Model 1.1 was then applied on images from this time and Model 1 to the remaining images of Magnus. Model 1 also transferred (i.e. served as a training starting point) on images of Mini and *M. lingua* within ROV video frames, producing Models 2 and 3 respectively (Figure 2). Model 2 was then applied to a GoogleDrive folder that contained all 9,173 images of Mini and Model 3 to a folder containing all 1,420 ROV video frames.

Model 4 was trained to identify the lasers in the same 1,420, 1400 x 888 pixel cropped ROV frames from the Tisler reef, independently of all other models. Inter-observer variation was removed as the same individual completed the training of all models.

### Stopping Criteria

Two distinct approaches were used to determine the endpoint of model training. Training cessation was guided by qualitative criteria for Models 1-3, but quantitative criteria only for Model 4. Training was deemed complete for Models 1 and 2 (LoVe Observatory) when predictions for at least two images from each month of the training dataset had required no corrective annotation. For Model 3 (ROV Tisler) segmentations that did not require any corrective annotation had to be seen for three frames from each video section; this more stringent criterion reflects the higher variability of image content and quality in the Tisler dataset. For Model 4 the simplicity of the subject of interest and its stark contrast to any background objects permitted the decision to stop training to be solely determined through RootPainter’s metrics calculations; specifically, when the rolling average (n=10) for the dice score reached 0.95.

The success of all models was also quantitatively assessed in real-time using the metrics calculated continually by RootPainter. Precision, recall, dice score, accuracy and estimated area error were exported to evaluate the suitability of this metrics calculator as stopping criteria in future work.

### Post-processing

Application of trained models to their respective datasets, via the ‘segment folder’ function in RootPainter, produced foreground predictions for every image. The discrete area values of each foreground prediction were extracted using the RootPainter ‘extract region properties’ function and exported as one .csv file.

The results of Model 1 and 2 (LoVe Observatory dataset) and Model 4 (Tisler reef dataset) were visually checked for anomalies. This involved scanning through the segmentation output file thumbnails for obvious errors, such as camera malfunctions, missing sponge/laser areas or obstructions by fish. This facilitated faster and more comprehensive datapoint exclusion than attempting to identify anomalies through pre-processing; post-processing required approximately 30 minutes of active work per 744 images analysed. Significant variation in sponge area and distribution in Tisler reef video frames prevented identification of Model 3 errors through segmentation observation alone. Given that additional approaches would suffer diminishing returns for an increase in result accuracy with user time, no post-processing was conducted for results from Model 3.

### Image Scaling

#### Images from the LoVe Observatory

In the absence of laser scales, the average width of a *Lophelia pertusa* branch from the Bømla reef in Hardangerfjord (0.43 cm ± 0.09, [15, Greiffenhagen et al. in prep]), and branches adjacent to Magnus and Mini were used to scale the pixel dimensions of images in ImageJ. This allowed conversion of the foreground areas predicted by RootPainter from pixels to cm^2^. Relative sponge areas were calculated through division of each surface area value by the maximum sponge area value for that dataset.

#### ROV Frames from the Tisler Reef

The x,y coordinates of areas segmented by Model 4 allowed calculation of the distance between laser points in each ROV video frame in pixels (Equation 1). Images were then independently scaled based on the true distance between the lasers, which is 5 cm. The area errors for each image, as calculated by RootPainter during training, were scaled in the same manner.

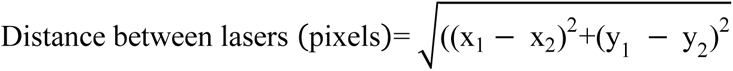

Equation 1: Formula used to calculate the Euclidean distance between laser points given their x,y coordinates, where (x_1_,y_1_) and (x_2_,y_2_) correspond to each laser point respectively.

### Model Validation and Statistical Analysis

Model 1 was validated by comparing surface area measurements made manually in Photoshop [48] and predicted by RootPainter for 452 randomly selected images (5% of the total dataset), 28 of which were seen during training. Sponge areas were extracted from the Photoshop annotations using open-source R scripts [48], and scaled using ImageJ as previously described. The precision, recall, dice score and accuracy of Model 1 were then calculated in Python by assuming the manually annotated images were accurate.

Precision is quantified as the ratio of true positives to all positive instances (the sum of true and false positives) and describes the probability that a pixel is truly foreground, given that the RootPainter model predicts it as foreground. Recall is calculated as the ratio of true positives to all true positive instances (the sum of true positives and false negatives) giving a measure of the proportion of foreground pixels the RootPainter model is expected to identify [49,50]. Dice score is calculated using precision and recall, giving an overall indication of model performance. Accuracy evaluates how close to the true result is to the model’s predictions based on the degree of overlap between predicted segmentations and the true regions [50]. In previous studies models have been defined as successful with a precision ≥ 0.71, recall ≥ 0.75, dice score ≥ 0.74 and accuracy ≥ 0.76 [4,9,26,51].

RootPainter also continually calculates these metrics during training but through assumption that corrected segmentations are accurate. Comparison of the corrective annotation metrics for Model 1 to the externally calculated validation metrics allowed evaluation of the necessity of separate manual validation for future RootPainter studies. Additionally, RootPainter provides estimates of area error during training allowing assessment of model success. Error is calculated through subtraction of the ‘corrected area’ from the ‘predicted area’ for each training image, where the corrected area is the post-annotation result, and is taken to be the true area. For Model 3 the agreement between the corrected area and RootPainter’s training prediction was also investigated through calculation of a Pearson correlation coefficient and linear regression, to ensure lack of bias across multiple sponge individuals of varying size.

## RESULTS

### RootPainter Model Development

In total, three models were produced and used to evaluate the surface area of *Mycale lingua*; a fourth model was produced to identify red lasers in ROV video frames. Table 1 displays the number of images and time used in both training and application of the models.

**Table 1:**
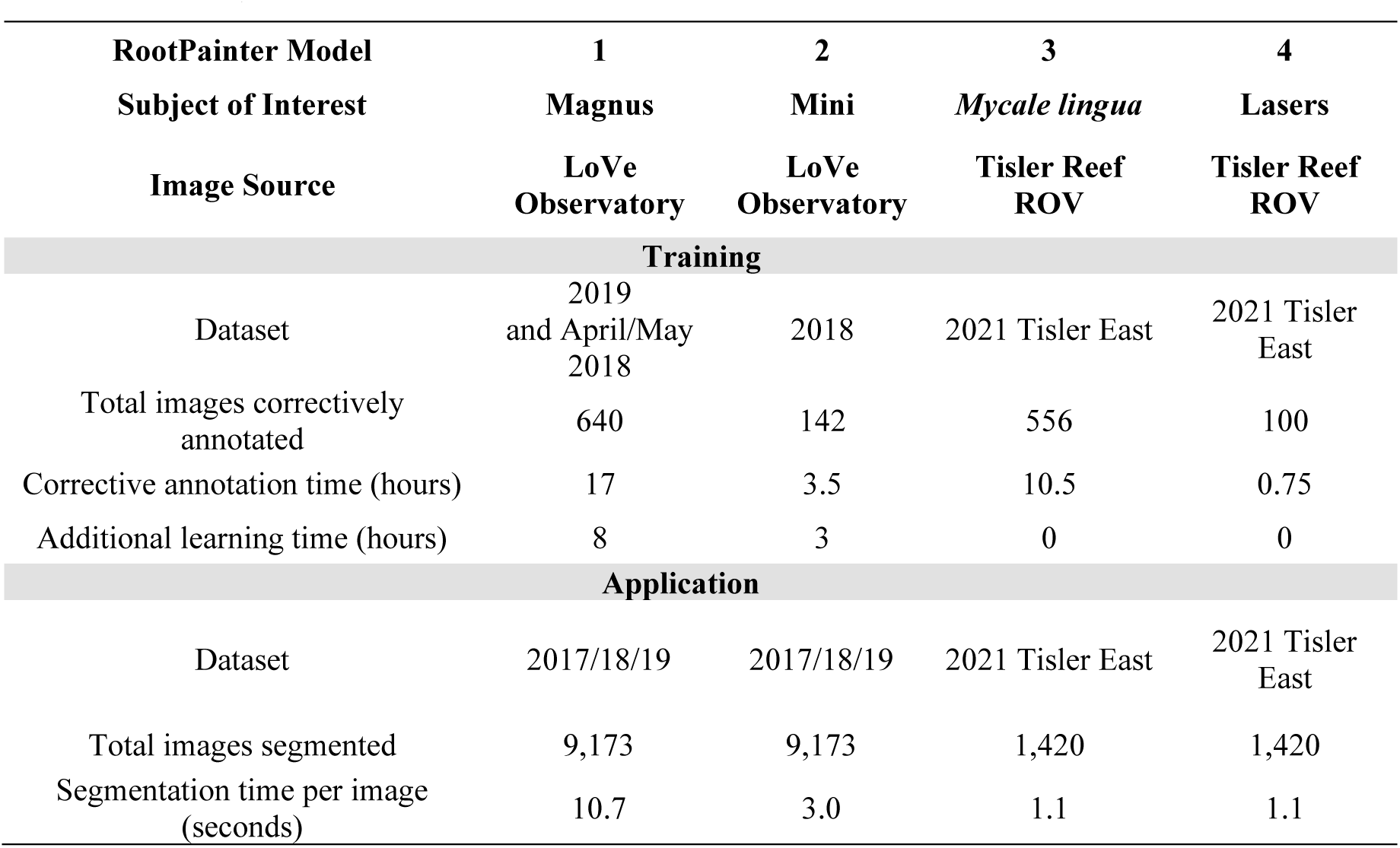
Training and application data for RootPainter Models 1, 2, 3 and 4. Additional learning time refers to time connected to GPU where no annotations were performed but training was left running to enable the model to better fit the existing annotations. The total images for Magnus include 120 additional images used to optimise Model 1 to turbid images during a colour texture change in April/May (Figure 2, Supplementary Information Table 1).

### Model 1

Model 1 was trained on 640 images of Magnus from the LoVe Observatory, requiring 17 hours. The training times for Model 1.1 (Supplementary Information Table 1), necessitated by the colour/texture change in Magnus during April and May of 2018/19, are incorporated into Model 1 in Table 1. The decision to stop training Model 1 was guided by qualitative criteria, but concurrent increases in the corrective annotation metrics of precision, recall, dice score and accuracy can be seen with improved segmentations in Figure 3.

**Figure 3:**
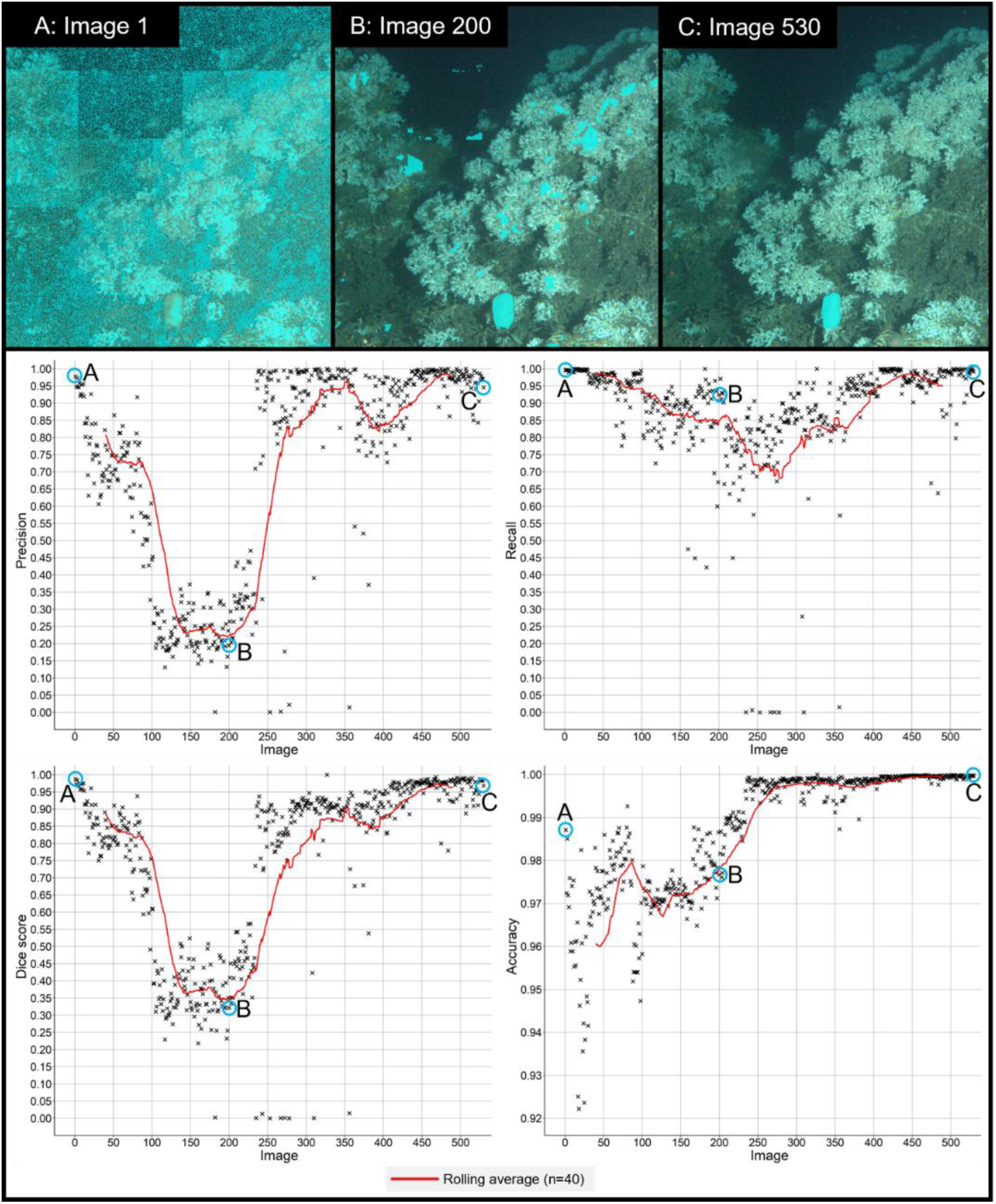
RootPainter predictions for images that appeared 1^st^, 200^th^ and 530^th^ during training of Model 1, where segmentation by the model is shown in light blue overlaying the input image. Accompanying corrective annotation metrics graphs displaying changes in precision, recall, dice score and accuracy of Model 1, as calculated by RootPainter during training. Values are displayed until image 530; the additional 120 images used in training RootPainter to recognise Magnus (Table 1), developed Model 1.1 (Supplementary Information Figure 1 and 2).

Model 1 was applied to 9,173 images of Magnus. Post-processing to exclude anomalies was completed, and highlighted that segmentations were impacted during March 2019, when sea-stars (suspected *Henricia* spp.) took prolonged residence on the base of Magnus. The area of sponge covered by the sea-stars varied, preventing reliable data point exclusion. Thus, segmentations from this period should be interpreted with caution. In total 548 data points were excluded, with 352 of these corresponding to corrupted images.

Figure 4 visualises the agreement between the areas of Magnus extracted using Model 1 and those manually measured in Photoshop. The average difference in area values between the methods is 2.26 cm^2^ ± 1.69 cm^2^ or 5.3% ± 3% of Magnus.

**Figure 4:**
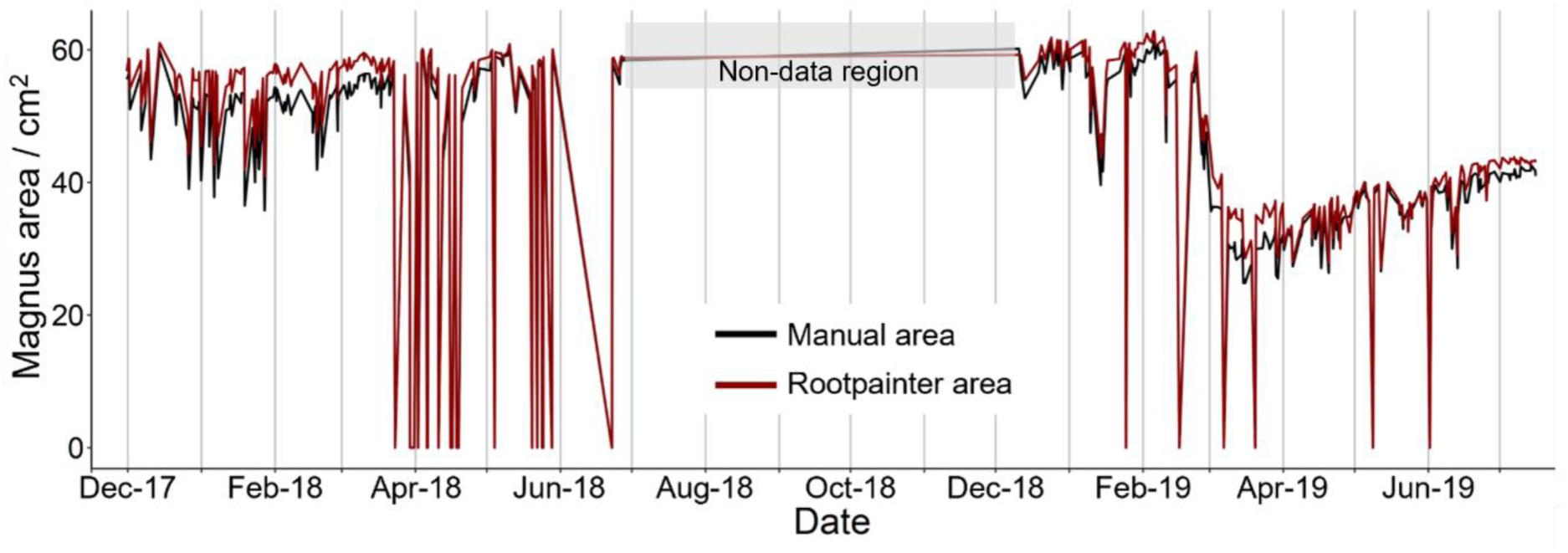
Comparison of Magnus’ area values as predicted by RootPainter and measured manually in Photoshop. Area highlighted in grey represents period during which no image data was available from the LoVe Observatory.

### Model 2

Fine-tuning of Model 1, through further training, was needed to produce Model 2 due to differential lighting of Magnus and Mini at the LoVe Observatory. This required an additional 142 images and 3.5 hours of corrective annotation on images of Mini, with the decision to stop training guided by qualitative criteria. The corrective annotation metrics from Model 2 training can be seen in Supplementary Information Figure 3. Model 2 was applied to 9,173 images of Mini, and post-processing completed to identify anomalies. In total, 601 data points were excluded; 352 of these corresponded to corrupted images.

### Model 3

Fine-tuning of Model 1 to produce Model 3 was necessary due to the more complex and changing nature of ROV video frames compared to underwater observatory images. This required 10.5 hours of corrective annotation on 556 video frames from the East of the Tisler reef, captured in 2021. The decision to stop training was guided by qualitative criteria, but the agreement between visual observations and the RootPainter corrective annotation metrics for Model 3 is demonstrated in Figure 5. Model 3 can distinguish *M. lingua* from *L. pertusa* (Figure 5B-C) and the sponge *Geodia* spp. (Figure 5A and C).

**Figure 5:**
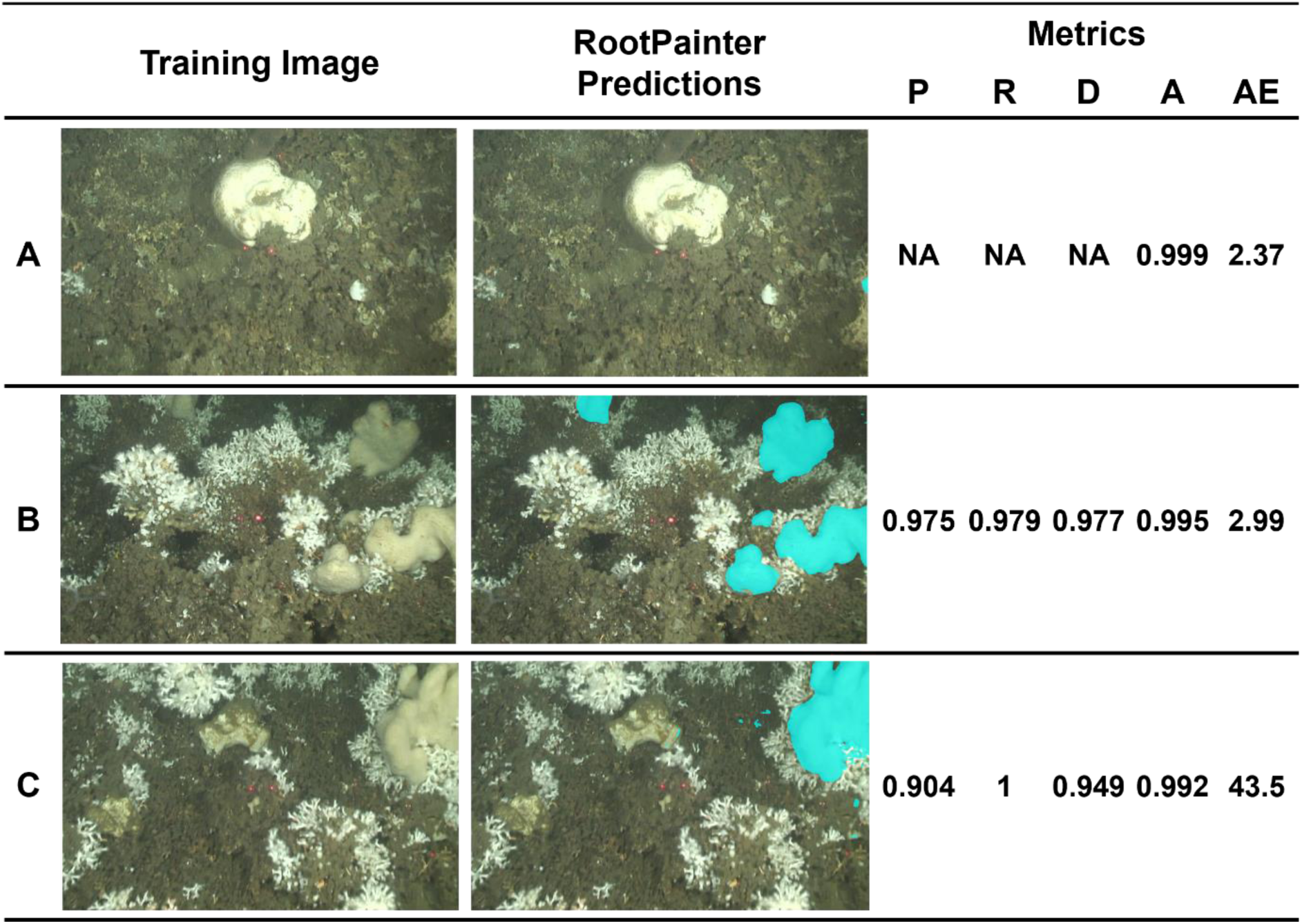
Examples of successful segmentations by RootPainter Model 3 and their accompanying metrics. Where P = precision, R = recall, D = dice score, A = accuracy and AE = area error / cm^2^. The images show; (A) *Geodia* spp. that is not mis-identified as *M. lingua*, (B) *M. lingua* individuals accurately segmented within *L. pertusa* and (C) *M. lingua* segmented accurately with nearby *Geodia* spp..

Model 3 was applied to all 1,420 ROV video frames from the East of the Tisler reef, captured in 2021. No post-processing was completed on the results from Model 3.

### Model 4

Model 4 was developed to segment red ROV lasers. It was trained on 100 video frames from the East of the Tisler reef, captured in 2021, requiring 45 minutes. The termination of training was solely determined by RootPainter’s metrics calculations (Supplementary Information Figure 4). Model 4 was applied to all 1,420 ROV video frames from the East of the Tisler reef, captured in 2021. Post-processing resulted in exclusion of 124 data points where only one laser was present.

### RootPainter Model Performance

#### Efficiency

RootPainter was 5-16 times more efficient compared to manual annotations (Table 2). Using RootPainter to analyse an ROV dataset requiring multiple annotations per image was more efficient than manual annotation of an underwater observatory dataset containing one individual per image (Magnus).

**Table 2:** Analysis time in seconds per image for Manual and RootPainter methods. Active user time for the manual method only includes the annotation times; for RootPainter it is the corrective annotation times and post-processing times combined. Inactive time for the manual method only includes the image area extraction time in R; for RootPainter this is the additional learning time and segmentation times (area extraction times were negligible for RootPainter). Magnus and Mini are both *M. lingua* individuals.

| Subject of Interest | Annotation Method | Image Source | Active user time (s per image analysed) | Inactive user time (s per image analysed) | Total time required (s per image analysed) |
| --- | --- | --- | --- | --- | --- |
| Magnus | Manual | Underwater Observatory | 105 | 26.0 | 131 |
| Magnus | RootPainter | Underwater Observatory | 9.09 | 13.8 | 22.9 |
| Mini | RootPainter | Underwater Observatory | 3.80 | 4.18 | 7.98 |
| *M. lingua* | RootPainter | ROV | 26.7 | 1.1 | 27.7 |

#### Accuracy

The precision, recall, dice score and accuracy for Model 1 is displayed in Table 3; agreement between the metrics as calculated by external manual validation and internal training calculations in RootPainter can be seen. Average corrective annotation metrics from the end-point of training Models 2-4 can be seen in Supplementary Information Table 3.

**Table 3:** Performance metrics for Model 1. Values from manual validation were calculated as a total result of overlaying all 452 manual annotations and their corresponding RootPainter predictions, meaning calculation of a standard deviation is not possible.

| Model | Calculation source | Precision | Recall | Dice Score | Accuracy | Training images used to calculate average |
| --- | --- | --- | --- | --- | --- | --- |
| 1 | Manual validation | 0.95 | 0.92 | 0.94 | 1.00 | NA |
| 1 | RootPainter corrective metrics | $0.97 \pm 0.04$ | $0.96 \pm 0.06$ | $0.96 \pm 0.03$ | $1.00 \pm 0.00$ | 430-530 |

#### Assessment of Model Success

Precision, recall, dice score and accuracy can reflect disproportionately harshly on model performance when foreground pixels are low (Supplementary Information Figure 5 and 6). These corrective annotation metrics were therefore used in combination with the area errors as calculated by RootPainter to assess the success of Models 1-3 (Figure 6). The agreement between the corrected/true area in each training image and RootPainter’s training prediction was also assessed for Model 3, as individuals of varying sizes are present in the ROV video frames (Figure 6d). For the final 400 images used in training the Pearson correlation coefficient between the corrected area and predicted RootPainter area is 0.95 (p-value < 2.2 x10^-6^); Model 3 consistently over-predicts the area of *M. lingua* by 4.89 cm^2^ as calculated by linear regression, with an R^2^ of 0.91.

**Figure 6:**
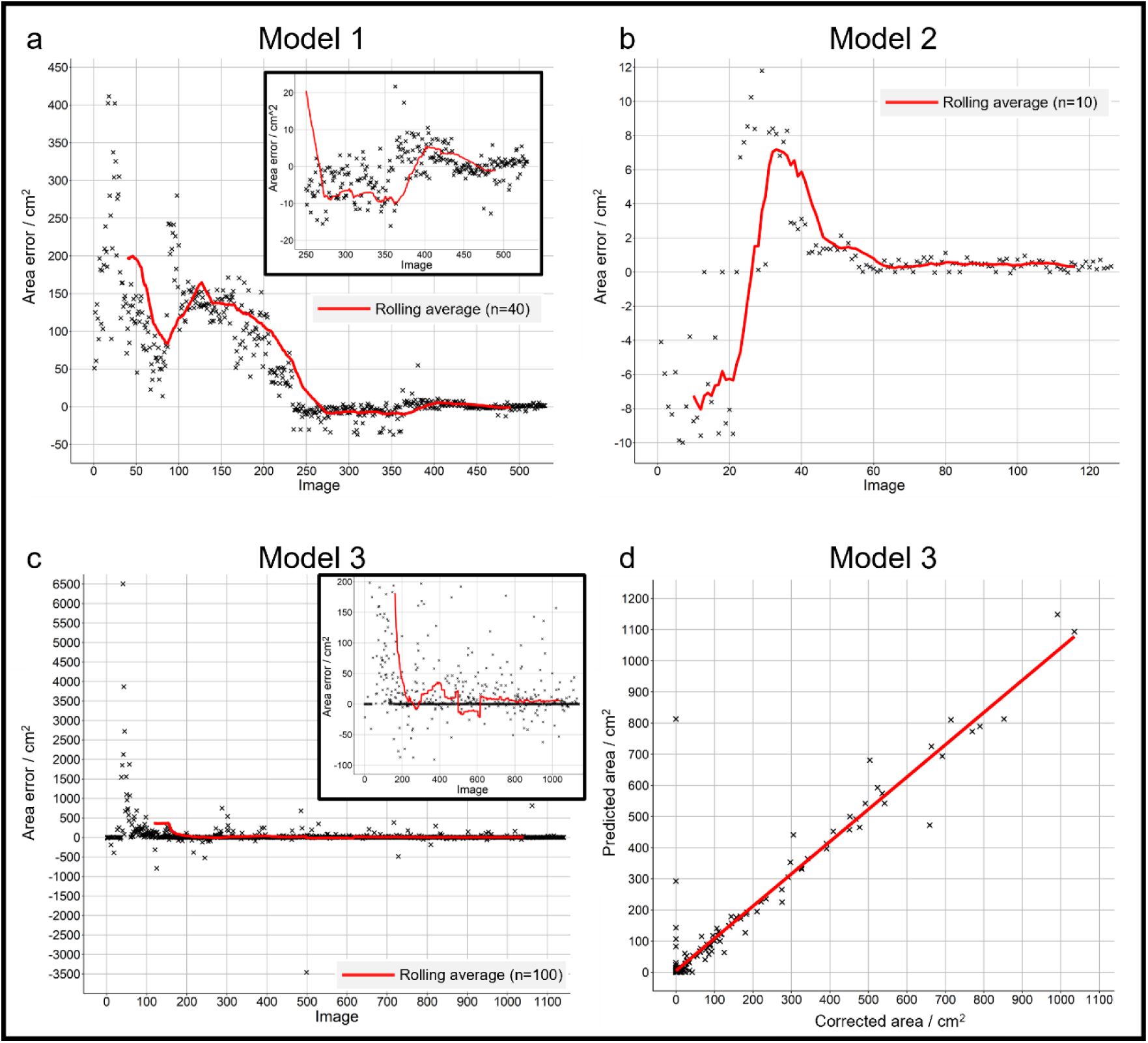
Graphs displaying changes in scaled area errors during training of: (a) Model 1, (b) Model 2 and (c) Model 3. (d) Graph demonstrating the correlation between *M. lingua* surface area as predicted by RootPainter and corrected during training for images 730-1130. The predicted area consists of all pixels RootPainter classified as *M. lingua* for each training image. The corrected area consists of all the pixels RootPainter classified as *M. lingua*, minus those the user highlights in green and plus additional pixels the user highlights in red.

The average area errors for each model, as calculated by RootPainter, towards the end of training can be seen in Table 4. The value for Model 1 is in agreement with the average area error calculated from manual validation (2.26 cm^2^ ± 1.69 cm^2^, Figure 4).

**Table 4:** Average area errors for Models 1, 2, and 3 as calculated by RootPainter during training. Average area error as a percentage was calculated using the average size of Magnus in 2019 for Model 1, Mini in 2018 for Model 2 as this was the data used in training the models. The percentage area error cannot be accurately estimated for Model 3 due to the wide range of sponge sizes within the data.

| Model | Average area error |  | Training images used to calculate average |
| --- | --- | --- | --- |
|  | cm<sup>2</sup> | % |  |
| 1 | $-0.06 \pm 2.87$ | $0.14 \pm 6.67$ | 430-530 |
| 2 | $0.45 \pm 0.86$ | $4.05 \pm 7.75$ | 100-150 |
| 3 | $7.09 \pm 52.97$ | NA | 730-1130 |

### Model Outputs and Observations

In total, 4 measurements were simultaneously extracted by RootPainter from the output segmentations of Models 1-3, including the area of individuals, as well as the diameter, perimeter and x,y coordinates of each discrete area. For the purposes of this study, we focused on the surface area outputs only.

In the LoVe Observatory dataset, 100% of the images contained the target species, *Mycale lingua*. The average 2D surface area for Magnus and Mini in the monitored months of 2018/19 are displayed in Table 5. In the Tisler reef ROV dataset only 40% of the extracted video frames contained *M. lingua* individuals, with an average size of 19.4 cm^2^ ± 51.8 cm^2^.

**Table 5:**
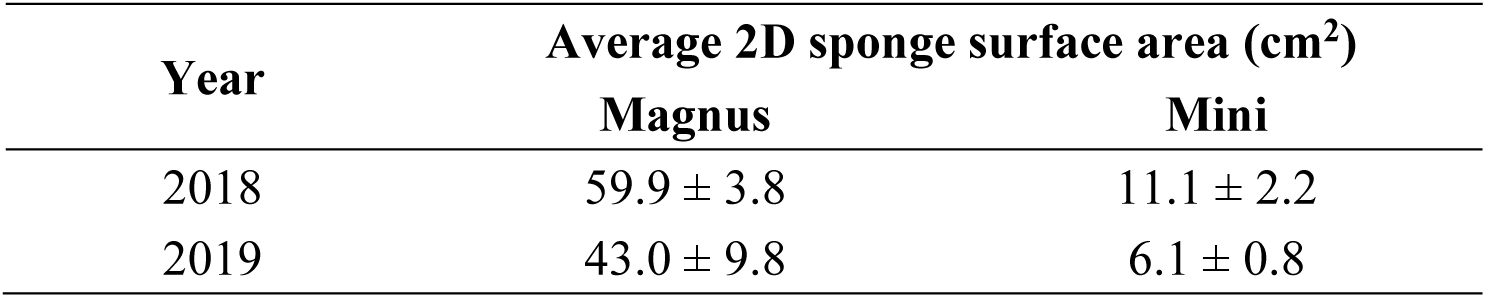
Average 2D size of Magnus and Mini at the LoVe Observatory in the monitored months of 2018/19.

| Year | Average 2D sponge surface area (cm<sup>2</sup>) |  |
| --- | --- | --- |
|  | Magnus | Mini |
| 2018 | $59.9 \pm 3.8$ | $11.1 \pm 2.2$ |
| 2019 | $43.0 \pm 9.8$ | $6.1 \pm 0.8$ |

Magnus and Mini both exhibited frequent contractions in each month of recorded data, without any displayed seasonality in this behaviour. A clear decrease in sponge surface area (∼50%) in the results from Model 1 compared to Model 2 was seen during February-March of 2019. Returning to the raw data revealed that this resulted from prolonged sea-star (*Henricia* spp.) residence and presumed predation at the base of Magnus. Mini’s area was unaffected during this time (Figure 7).

**Figure 7:**
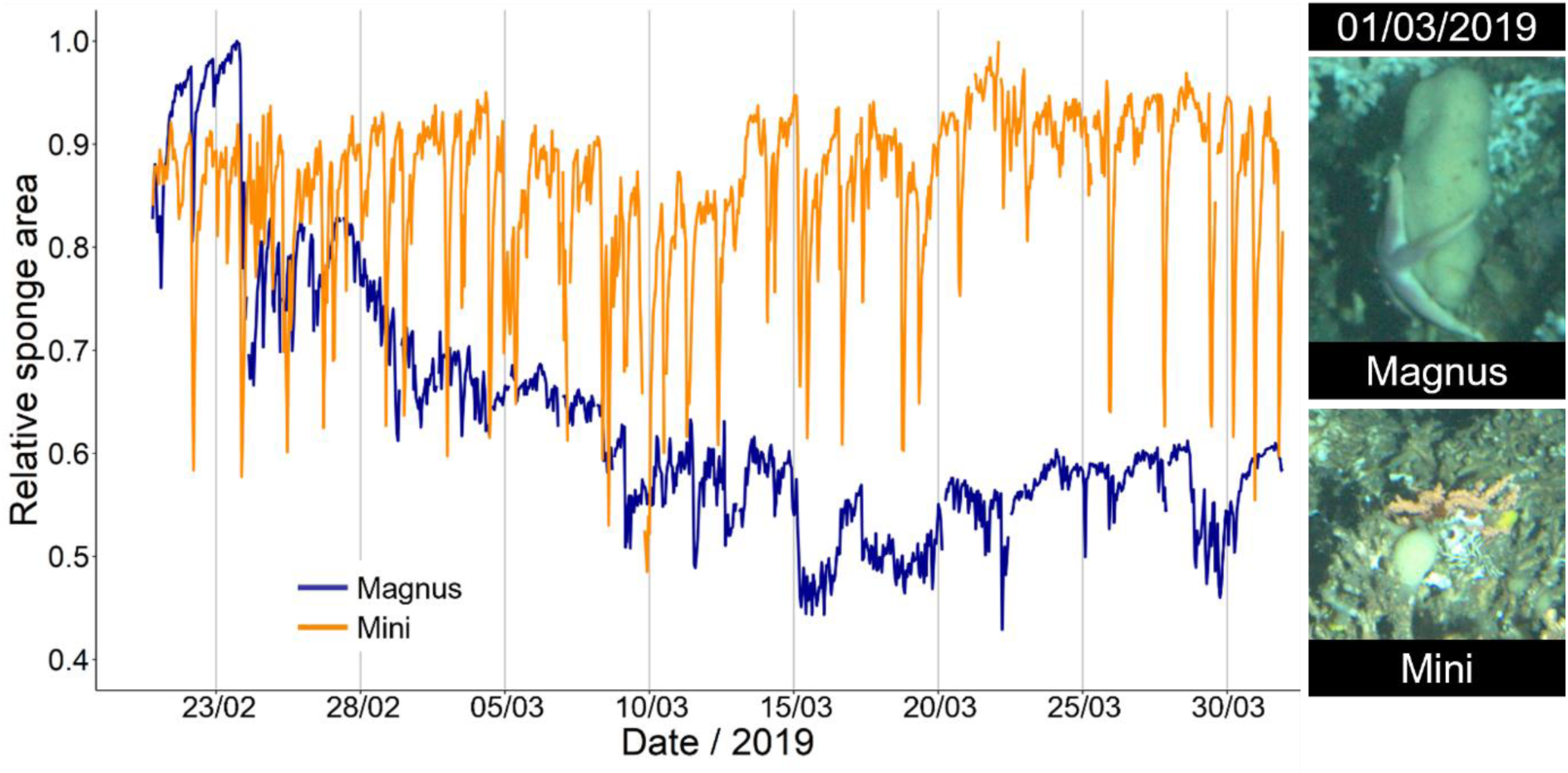
Relative areas of Magnus and Mini whilst sea-stars reside on the base of Magnus.

## DISCUSSION

This study showed the suitability of the user-friendly machine learning tool, RootPainter, to analyse large datasets of marine images. RootPainter was capable of accurately processing images of varying size, colour, and complexity, five to sixteen times faster than manual annotation, without the need for image pre-processing. As the efficiency of RootPainter is dependent on dataset size this may be even faster for larger datasets. Manual validation demonstrated the reliability of our qualitative stopping criteria and RootPainter’s in-built metrics calculator as a means to assess model success. Therefore, external stopping criteria and model validation may not be required in future studies, allowing ecological conclusions to be drawn, with the appropriate caveats in place, with significantly improved efficiency.

### Machine Learning Tools for Marine Image Analysis

This work demonstrates that RootPainter is an accessible and affordable tool capable of processing large and complex datasets, with the potential to ease the analysis bottleneck created by the continually increasing volume of video/image data collected by marine researchers. The intuitive interface and instruction notebook accompanying the software allow marine experts to concentrate on dataset content, instead of the intricacies of running a machine learning algorithm. This includes removing the need for image ‘pre-processing’ steps, such as reduction of background complexity, as seen for other machine learning methods. The ability to run RootPainter through GoogleColab prevents users needing to acquire an expensive GPU or to possess significant computing power. In order to comply with the free GoogleColab GPU usage and GoogleDrive space limits, annotations in this study were completed in 3-5 hour sessions and images were uploaded in batches. Consequently, powerful results were produced with no previous user experience in machine learning and at no additional cost.

This study focused on RootPainter’s application to *M. lingua* individuals only. However, the success demonstrated with this notoriously complex species of interest, paired with previous terrestrial examples of model success [31,52–54], gives confidence that RootPainter will be capable of segmenting other marine species. The applicability of RootPainter to species identification and biodiversity investigations may be increased by introduction of multi-annotation capabilities. Within the current version of RootPainter simultaneous investigation of multiple species requires the development of several different models for each target species (i.e. class) and extraction of results separately. Alternatively, a multi-staged approach can be used where the general foreground is segmented first and then this is used to remove all background from the data. The extracted foreground could then be further categorised into different classes. Whilst this model-cascade approach may have training and efficiency benefits, akin to localisation [55], its utility decreases with increasing class number. Therefore, depending on user needs, a conscious decision regarding choice of machine learning tool for a desired investigation needs to be made.

The web-based annotation software BIIGLE is widely used by marine ecologists for manual annotations and is capable of automated novelty detection [56]. The user friendly and open-source ‘machine learning assisted image annotation’ (MAIA) function in BIIGLE has proven suited to biodiversity studies due to its multiclass annotation capabilities [25]. However, compared to RootPainter, and at the time of writing (October 2023), extraction of information such as species area or perimeter cannot be automated in BIIGLE, no training metrics are provided to aid assessment of model success, and models developed through the MAIA function cannot currently be transferred between users through the existing interface.

ImageJ is also widely used for manual analysis of marine images. Whilst it does not possess its own machine learning tool as such, the deepImageJ plugin enables users to apply pre-trained neural networks (models) in ImageJ, that are downloadable from an ‘online zoo’ [57]. Application of downloaded models is user-friendly, but their *de novo* development or continued training must be completed outside the ImageJ application [58]. This requires machine learning expertise, creating dependence of non-experienced users on others to develop models they require. However, as deepImageJ users are downloading ‘bespoke’ algorithms, the range of functionalities models can possess and the information that can be extracted from images is expansive. Thus far deepImageJ has been targeted at microscopy work and biomedical imaging, such as virtual tissue staining [59] and instance segmentation of neurons [60]. When analysing marine images, the inability to optimise models without computational expertise would act as a significant barrier to the use of deepImageJ, as the quality and background of underwater images varies significantly. The accessibility of model sharing and application within deepImageJ has undoubtedly made significant progress to unifying the field of microscopy image analysis [57], with some models being downloaded 20,000 times [58]. The additional flexibility provided by RootPainter’s accessible training process may enable a similar achievement to be made in the marine imaging field.

### RootPainter Model Sharing

Sharing RootPainter models may allow researchers to dramatically increase their marine image analysis capacity and analyse datasets to their full potential. However, the time-saving capabilities of transferred models within RootPainter likely depends on the specific task and datasets utilised [61]. Starting training with a suitable pre-established model may reduce the time and number of images required to produce a satisfactory model for a given dataset; only 3.5 hours and 142 images, and 10.5 hours and 556 images were required to optimise Model 1 to produce Models 2 and 3, respectively (Table 1). The initial increased accuracy of RootPainter predictions, as seen for the first 20 images of Model 2 compared to Model 1 (Figure 3 and Supplementary Information Figure 4), significantly reduces the corrective annotation time required per image (Table 2). Pre-developed models can also reduce the threshold number of ‘application images’ at which using RootPainter becomes more efficient than manual annotation. For *de novo* model development on static observatory images a minimum of 468 images are needed in the application dataset to ‘justify’ use of RootPainter. Utilising a pre-developed model reduces this to 96 static observatory images or 289 ROV video frames (Table 1 and 2). However, greater image numbers may be required depending on the dataset (Table 1).

The accuracy of transferred models will always be limited by object variation between datasets. Researchers are therefore advised to utilise the adaptability of the RootPainter algorithm to fine-tune a pre-developed model to their dataset before its application. This will also produce corrective annotation metrics, allowing users to assess the success of segmentations themselves.

### Analysing Static vs Mobile Image Datasets with RootPainter

#### Dataset effect on speed of training

Image analysis with RootPainter is highly efficient for both static images and frames from moving videos (Table 2). However, the number of images and training time required for model development on a given species does increase when moving from underwater observatory images (Models 1 and 2) to ROV video frames (Model 3). The more dynamic background, reduced image clarity and varied lighting within the ROV video frames, as well as the need to identify and distinguish many different *M. lingua* individuals from apparently similar *Geodia* spp., increased the extent of model optimisation required to produce Model 3 compared to Model 2 (Table 1).

Interestingly, the rate of corrective annotation in RootPainter did not increase with increasing image complexity; optimisation of Models 2 and 3 did not involve significant background annotations, with both requiring 1.2 minutes of annotation per image (Table 1). Conversely, the rate of manual annotations does decrease with increasing image complexity (i.e. more individuals per image require more time to manually annotate). Therefore, comparison of the development speed of Model 3 to underwater observatory manual annotations likely underestimates the efficiency of RootPainter for ROV video frame analysis.

It is important to note that the nature of the subject of interest also impacts RootPainter model training time. As the red lasers were uniform in each image and visually distinct from all other background objects development of Model 4 required the least time and number of images, despite being trained ROV video frames.

#### Dataset effect on accuracy of models

All RootPainter models in this study exhibited high levels of accuracy (Table 3, Supplementary Information Table 3). Manual validation confirmed that Model 1 consistently and accurately predicted the area of Magnus (Table 3, Figure 4). Poor agreement between the Photoshop and RootPainter results was often a result of external factors, such as sea-star presence or turbidity (Supplementary Information Figure 7). The average area error of Model 1, as calculated by manual validation, is larger (5.3% ± 3%) than the estimate provided during training by RootPainter (0.14% ± 6.67%), but not significantly so. Therefore, we may accept the average area error estimates for Models 2 and 3 calculated by RootPainter during training (0.45 cm^2^ ± 0.86 cm^2^ and 7.09 cm^2^ ± 52.97 cm^2^, respectively), to be representative of the true accuracy of area predictions for these models (Table 4).

The accuracy of Models 1-3 can be seen to decrease with increasing dataset complexity. Model 1 was trained on, and used to segment, images of the same individual. Conversely, each image segmented by Model 3 contained different sponge individuals, including many which the model was not exposed to during training. The effect of this is apparent in the larger average area error for Model 3 than for Model 1 (Table 4). However, it should be considered that the accuracy of manual annotations may also decrease across these two datasets, and the overall area error for Model 3 is still acceptably small.

### Increasing the Efficiency of RootPainter

The efficiency of image analysis with RootPainter depends on the images and computing set-up used. Without non-trivial pre-processing to reduce image complexity, the main methods to increase analysis efficiency involve using smaller images, paying for upgraded GoogleColab access, or investing in a purpose built deep-learning workstation. As the aim of this tool is to be accessible and cost-effective, use of equipment designed for deep-learning will not be discussed further here.

Smaller images increase the efficiency of image analysis with RootPainter through increased training speeds and reduced application times [31]. When constrained to larger images, using the ‘create dataset’ function in RootPainter to randomly crop training images can produce a more efficient training dataset [31]. Smaller images require less time to segment during model application; Model 2 was applied to images 2.8 times smaller than Model 1 (Figure 2), and they were segmented 3.6 times faster (Table 1). The same application segmentation speeds seen for Models 3 and 4 (Table 1), demonstrate that subject complexity does not affect RootPainter application time.

Upgrading GoogleColab can significantly reduce both the ‘active’ and ‘inactive’ user time required for RootPainter studies, through increased access to higher memory GPUs. Chance assignment to a higher memory GPU resulted in reduced segmentation times during application of Models 3 and 4 compared to Model 2 (Table 1), despite their application to images of similar sizes (Figure 2). The application efficiency of RootPainter may therefore be tripled if improved GPU assignments can be consistently secured through a paid upgrade in GoogleColab (∼£8 a month in the year 2023).

Finally, excluding the optional post-processing stage in this study would have reduced the total ‘active user time’ by 6.2 hours each for Models 1 and 2, increasing the efficiency of RootPainter to 6 and 25 times faster than manual annotation for these models, respectively (Table 2).

### Improving the Accuracy of RootPainter

Using smaller images may increase segmentation accuracy due to mitigation of class balance issues; large background to foreground ratios are known challenges for convolutional neural network model training [62]. This may be reflected in the smaller standard deviation for average area error of Mini as predicted by Model 2 within smaller images, than for Magnus (−0.06 cm^2^ ± 2.87 cm^2^) as predicted by Model 1 (0.45 cm^2^ ± 0.86 cm^2^).

The post-processing (i.e. exclusion of obvious segmentation anomalies) completed in this study aimed to improve the accuracy of results from Models 1 and 2. Of the 452 images used in manual validation, 5 segmentations (including ‘16/04/2019 22:09’, Supplementary Information Figure 7) had been removed during post-processing of Model 1. This caused no improvement in the precision, recall, dice score and accuracy of Model 1, to two decimal places. Therefore, the post-processing stage may not be necessary in future studies.

Ultimately, the accuracy of a RootPainter model depends on the quality of user corrective annotations, and whether the training images are sufficiently representative of the subject of interest. Even manual annotations incur some error due to ambiguity on the boundary of subjects, the difficulty of perfect annotation and partial volume issues. This also results in diminishing returns in accuracy from continued annotation towards the end of model training (Figure 6). Therefore, accepting small inherent error in segmentations is essential to maintaining efficiency in RootPainter studies.

### Reliability of Metric Calculations in RootPainter

The corrective annotation metrics calculated within RootPainter during training overestimated precision, recall, dice score and accuracy by 0.02-0.04 compared to values from external manual validation for Model 1 (Table 3). Accounting for this overestimation the corrective metrics values for Models 2-4 still fall within the classification of successful models [4,9,26,29,51]. The discrepancy between calculations may result from images with regions of high uncertainty as during corrective annotation users can leave ambiguous errors as unclassified, conversely during manual annotation the user was forced to classify with certainty each pixel of an image. If this potential error is considered, utilising the corrective annotation metrics within RootPainter may negate the need for time-consuming manual validation in future studies. However, this decision should be left to users’ discretion, and it may be advised to complete manual validation when developing a model for a new species.

The overall reliability of corrective metrics calculations within RootPainter (Table 3) allows identification of when the user can stop training and accurate model performance is achieved. This was trialled to success with Model 4, thus providing a possible mechanism to reduce subjectivity in training cessation across RootPainter users. However, RootPainter’s metric calculations can be skewed by imperfect user corrections. For example, in the early stages of Model 1 training the extensive background pixels were not fully correctively annotated, to avoid overwhelming the algorithm, resulting in incredibly high metrics at a time where segmentations are poor (Figure 3). As Model 1 then improved its corrective metrics initially decreased as corrections became more thorough, before increasing again with the true accuracy of the model. Metrics may also be misleading for subjects of interest more complex than lasers (Supplementary Information Figures 5 and 6). Interestingly, the area error estimate by RootPainter continuously agrees with the visual assessments and can differentiate between good (Figure 5C) and excellent (Figure 5B) segmentations. RootPainter’s calculation of area errors would thus make an excellent contribution to stopping criteria in future studies investigating species area. Therefore, it is recommended that when complex subjects of interest are targeted, a combined qualitative and quantitative stopping criteria approach is used.

### RootPainter Applications

Machine learning tools for image analysis have the potential to rapidly increase our understanding of marine species and their functions within ecosystems. In this study, RootPainter has demonstrated an aptitude to identifying and predicting the surface area of *M. lingua*, both in underwater observatory images and ROV video frames. Due to the high ratio of background to foreground pixels in images used Models 1-3 slightly overestimated sponge area (Figure 4, Table 4). This error is very small and insignificant to the intended purposes of Models 1 and 2, investigating relative changes in the predicted sponge area for Magnus and Mini over time. Conversely to estimate sponge cover or biomass, as is the purpose of Model 3, a small and consistent error in predicted area is important. Whilst this is difficult to achieve with mobile ROV video frames, the average area error of 7.09 ± 52.97 cm^2^ for Model 3 is sufficiently small that ecological conclusions can be drawn if it is taken into consideration. The large standard deviation of this area error does not represent model bias disproportionately affecting larger or smaller sponges as there is a strong correlation between the user corrected and RootPainter predicted areas for Model 3 (Figure 6d). Finally, the ability of Model 3 to distinguish between apparently similar sponges, *M. lingua* and *Geodiia* spp. (Figure 5C), confirms that the model is reliable for investigation of ecological questions pertaining to a given species.

Results from Models 1 and 2 demonstrated the utility of RootPainter to temporal changes in species behaviour. Sponge contractions have previously been studied in both shallow and deep-water using manual and bespoke machine learning methods [2,21,28,29]. ‘Intrinsic’ contractions observed in shallow-water sponges likely serve to clear the aquiferous system, where blocked canals may disrupt filter-feeding [63,64]. In abyssal sponges the contracted state can be maintained for up to weeks at a time and is believed to represent an energy saving mechanism through reduction of sponge filter-feeding [2]. In this study, *M. lingua* exhibited short and frequent contractions consistently throughout the seasons, suggesting an alternate purpose for some sponge contractions to energy conservation and aquiferous system clearing is likely. Contractions were consistently ‘larger’ for Mini than for Magnus relative to their overall size, but contraction rate is similar between the sponges. In April 2019 prolonged sea-star residency on Magnus coincides with a ∼50% reduction in sponge size and a significant reduction in sponge contractions. This energy conservation may represent a viable survival strategy for during predation; contractions in Mini were unaffected during this time. Previous investigation of *M. lingua* contractions found them to be rare and asynchronous at 30 m depth [42], but frequent and correlated with salinity in one individual at 260 m depth [29]. The possibility that environmental drivers are contributing to the observed behaviour of Magnus and Mini at the LoVe Observatory requires further study. This may elucidate the purpose of the non-energy conservation contractions seen through identification of any environmental stimuli. Variation in contractions with abiotic factors will have implications for the ecosystem services provided by deep-sea sponges, especially if frequent contractions are concluded to effect filtration capacity.

The suitability of RootPainter to spatial analyses, such as investigations into species distributions, has been shown through successful development of Models 3 and 4. Quantifying deep-sea sponge presence and surface area allows estimation of their percentage cover and/or biomass, and therefore contribution to carbon-cycling in benthic environments [44,65]. The distribution of *M. lingua* has previously been investigated at the Tisler reef through small datasets [7,10], but determination of its variation in time and space across the reef has been prohibited by methodological limitations. Requiring just two working days, RootPainter produced results for the distribution, abundance, and size of *M. lingua* across the East of the Tisler reef. Thus, the use of machine learning tools, such as RootPainter, will be essential in the future study of spatiotemporal patterns within large image datasets.

## CONCLUSION

RootPainter provides a viable solution to the increasing data processing needs of marine ecologists, both on time-lapse data from static underwater observatories and frames from ROV/AUV video data. Through proper training the algorithm can efficiently produce highly accurate models, and its built-in methods to assess stopping criteria and model success reduce the need for manual validation. Additionally, regular improvements to the software continually enhance its suitability to marine image analysis; completion of the multi-annotation capabilities of RootPainter currently under development would increase the range of ecological questions that can be tackled using RootPainter. Resource limitation is not prohibitive to accessing this user-friendly software and the adaptability of models has the capability to productively link marine image analysis researchers. Moving forward the creation of a RootPainter repository to facilitate model sharing between users has the potential to exponentially increase the rate of information extraction from marine images, and therefore our understanding of marine organisms.

## AVAILABILITY OF DATA AND MATERIALS

All data used and models produced in this work are actively being uploaded to Pangaea.

## ACKNOWLEDGMENTS

We would like to thank the Lofoten Vesterålen Ocean Observatory, and specifically Geir Pedersen, for supplying much of the data utilised in this study.

H.P.C is supported by the Biotechnology and Biological Sciences Research Council EASTBIO Doctoral Training Programme (BB/M010996/1). A.G.S is supported by Novo Nordisk Foundation grant NNF22OC0080177. D.M.F is supported by the Rural and Environment Science and Analytical Services Division (SRUC-C5-1). L.D.C received funding from the European Union’s Horizon 2020 iAtlantic project (Grant Agreement No. 818123). This manuscript reflects the authors’ views alone and the European Union cannot be held responsible for any use that may be made of the information contained herein.

