## Supplementary Information for "New Interactive Machine Learning Tool for Marine Image Analysis"

### Model 1.1

Due to turbidity issues and changes in the colour/texture of Magnus during April/May of 2018/19, Model 1 required separate optimisation on these months, producing Model 1.1 as a result (SI Table 1).

SI Table 1: Training and application data for RootPainter Models 1, 2 and 3. Additional learning time refers to time connected to GPU where no annotations were performed but algorithm was left running so higher epochs could be reached.

| **Sponge** | **Magnus** | **Magnus** |
|---|---|---|
| **RootPainter Model** | **1** | **1.1** |
| **Training** | | |
| Dataset | 2019 | April/May 2018/19 |
| Total images used | 530 | 120 |
| Images annotated | 530 | 110 |
| Corrective annotation time / hours | 14.7 | 2.3 |
| Additional learning time / hours | 2 | 6 |
| **Application** | | |
| Dataset | 2017/18/19 April/May excluded | April/May 2018/19 |
| Total images segmented | 6345 | 2,828 |
| Segmentation time per image / seconds | 9.8 | 12.7 |

The same qualitative stopping criteria was used to determine the end point of training for Model 1.1 as for Model 1. The accompanying training metrics for Model 1.1 can be seen in SI Figure 1.

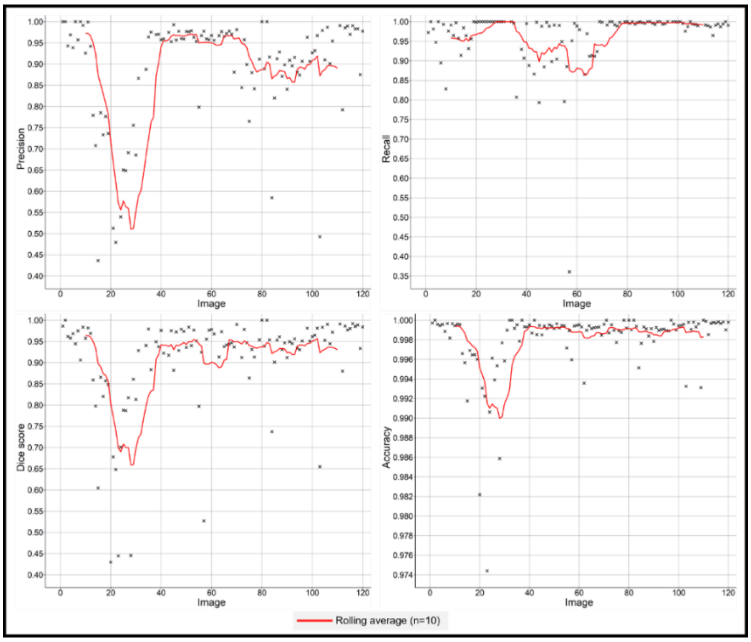


SI Figure 1: Graphs displaying changes in precision, recall, dice score and accuracy of Model 1.1, as calculated by RootPainter during training.

The average values for these metrics towards the end of training are displayed in SI Table 2. The precision for Model 1.1 is reasonably low. A low precision may result from many false positives or very few true positives being predicted. In this case it is likely the latter; the training images for Model 1.1 (April and May of 2018/19) were impacted by turbidity that often fully obscured Magnus. This combined with a small number of false positive pixels representing turbidity with ‘sponge-like’ colour or texture resulted in the lower precision for Model 1.1.

SI Table 2: Average RootPainter metrics for Model 1.1 calculated using the final 20 images/segmentations of training.

| Model | Calculation source | Precision | Recall | Dice Score | Accuracy |
|---|---|---|---|---|---|
| 1.1 | RootPainter training images 100-120 | 0.93 ± 0.12 | 0.99 ± 0.01 | 0.95 ± 0.08 | 1.00 ± 0.00 |

The change in area error for Magnus during training of Model 1.1 is shown in SI Figure 2. For the final 30 training images, the average area error in segmentations from Model 1.1 was 3.72 ± 8.52 $cm^2$. This is the highest area error and standard deviation seen out of the underwater observatory Models. Considering the nature of the dataset for Model 1.1, this is to be expected; the training images were taken from periods with high storm presence and resulting turbidity, as well as a physical appearance change in Magnus. Therefore, there is greater variation in the error of predicted areas during this time as a result.

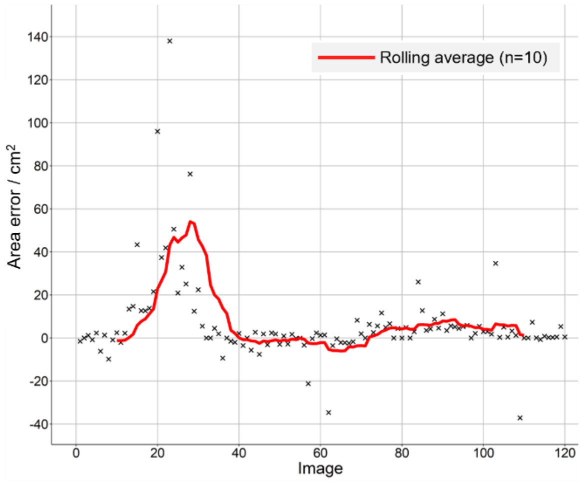


SI Figure 2: Graph displaying changes in scaled area errors during training of Model 1.1 on images of Magnus from April/May of 2018/19.

### Additional Corrective Annotation Metrics

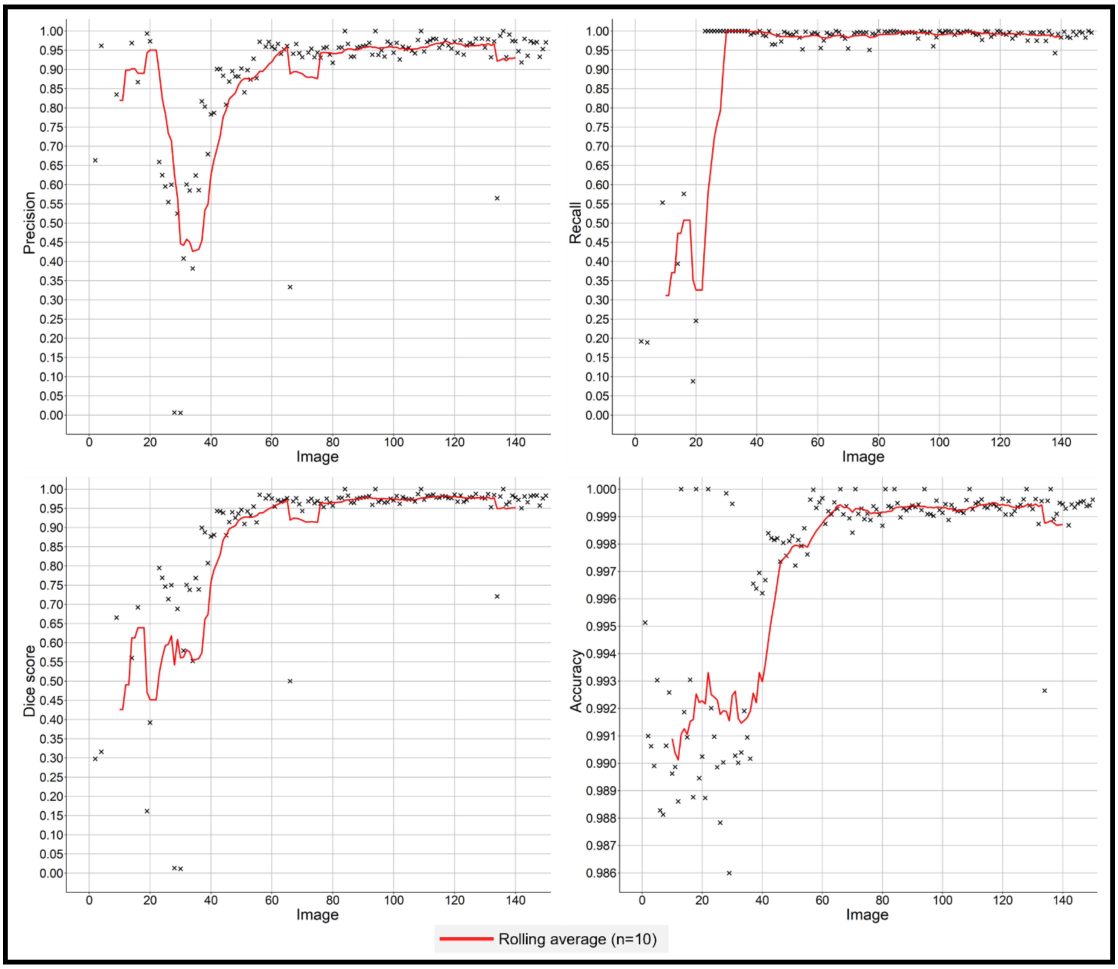


SI Figure 3: Graphs displaying changes in precision, recall, dice score and accuracy of Model 2, as calculated by RootPainter during training.

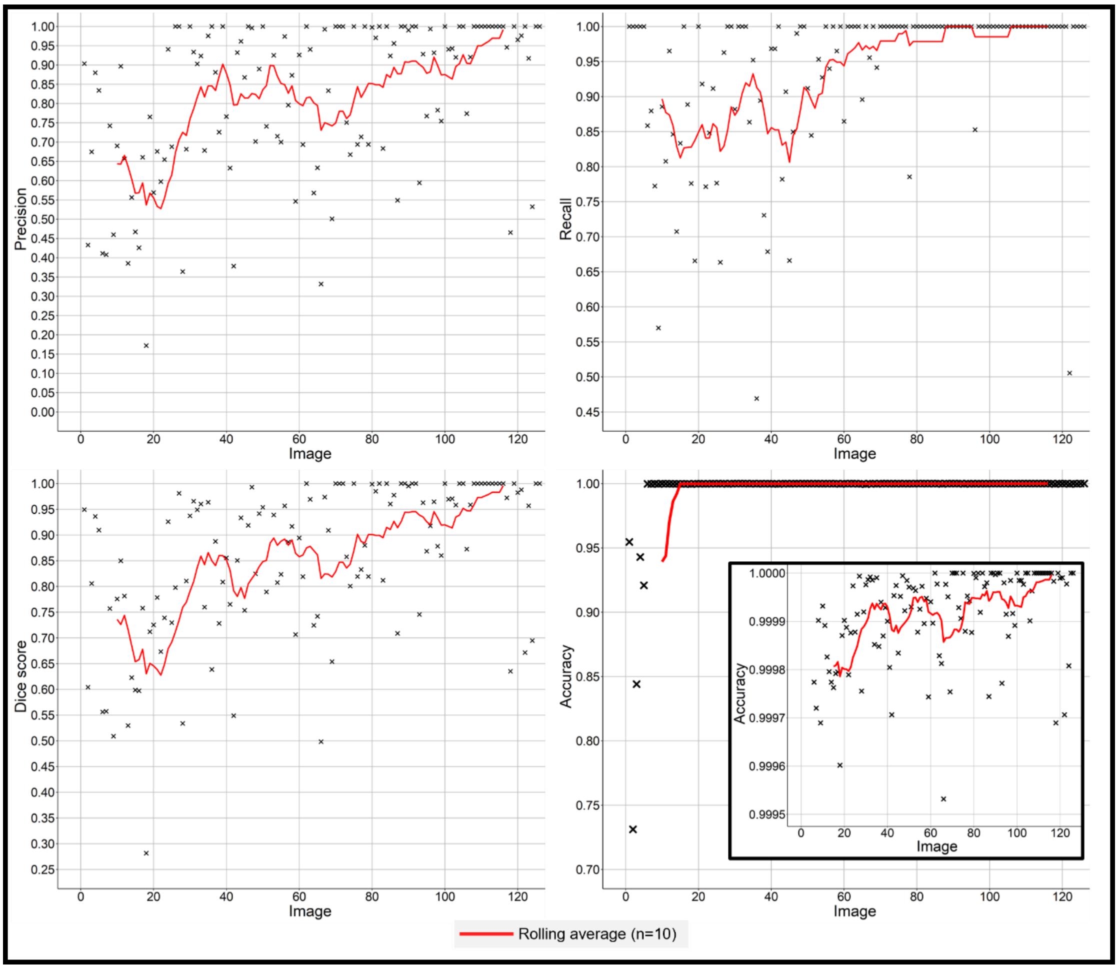


SI Figure 4: Graphs displaying changes in precision, recall, dice score and accuracy of Model 4, as calculated by RootPainter during training.

SI Table 3: Corrective annotation metrics for Models 2-4.

| Model | Calculation source | Precision | Recall | Dice Score | Accuracy | Training images used to calculate average |
|---|---|---|---|---|---|---|
| 2 | RootPainter | 0.95 ± 0.06 | 0.99 ± 0.01 | 0.97 ± 0.04 | 1.00 ± 0.00 | 100-150 |
| 3 | RootPainter | 0.87 ± 0.21 | 0.89 ± 0.22 | 0.84 ± 0.22 | 1.00 ± 0.01 | 730-1130 |
| 4 | RootPainter | 0.95 ± 0.12 | 1.00 ± 0.00 | 0.97 ± 0.08 | 1.00 ± 0.00 | 100-120 |

### Assessing Success of Model 3

Corrective annotation metrics can reflect disproportionately harshly on the success of a model. Therefore, they were used in corroboration with visual assessments to determine how successful Model 3 was. For example, the corrective annotation metrics for Model 3 are displayed in SI Figure 5, and the segmentations responsible for the highlighted ‘poor metrics’ in SI Figure 5 are shown in SI Figure 6; for A the area of *M. lingua* has only been over-estimated by 4.68 $cm^2$, but the precision, recall and dice score are all below 0.02.

In SI Figure 6A, C, D and E, the segmentations from training Model 3 appear as acceptable, but report very low precision, recall, dice scores and accuracies. Conversely, the metrics for SI Figure 6B, F and G, are generally improved, but upon inspection the segmentations would be deemed poor.

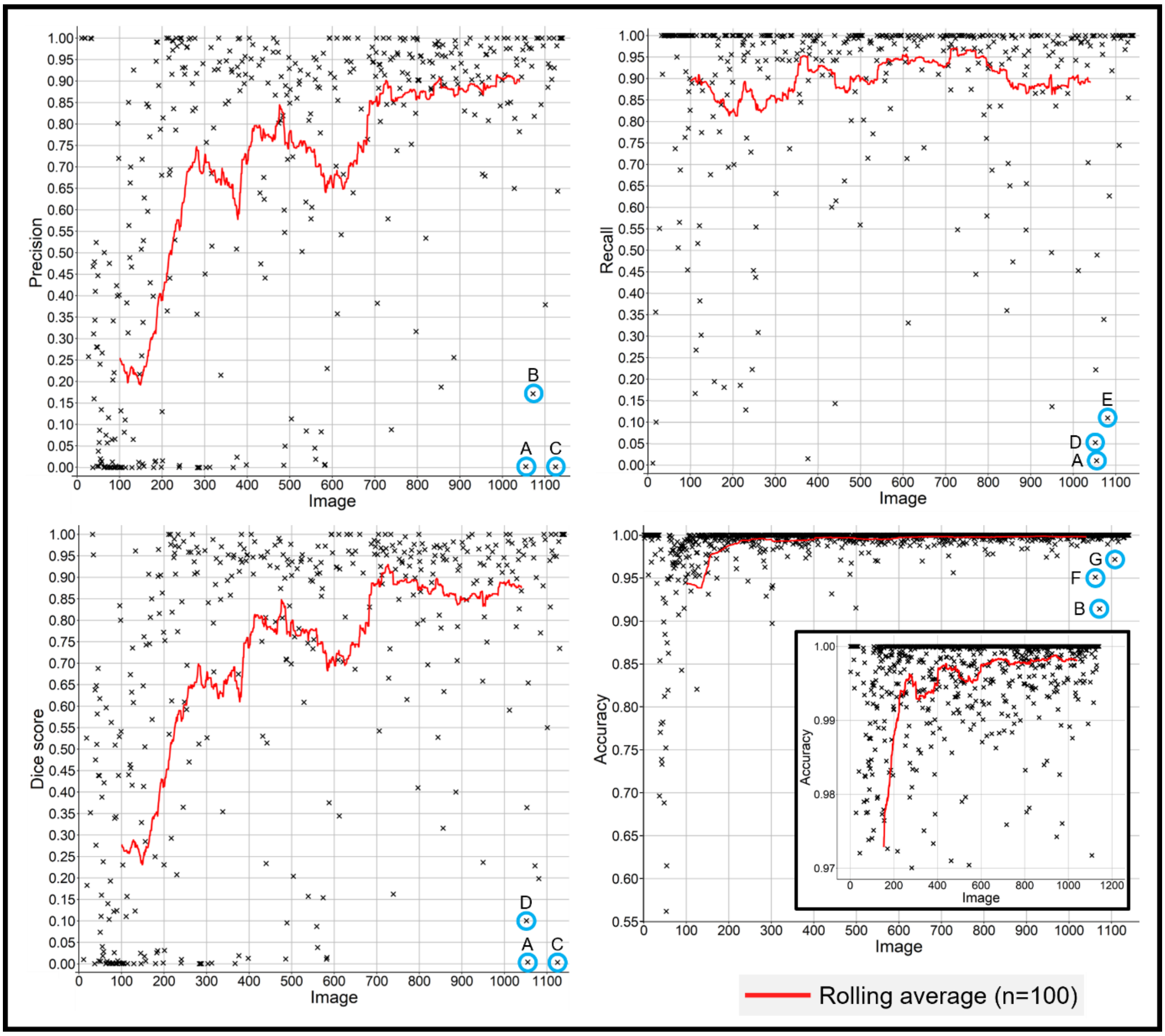


SI Figure 5: Performance metrics graphs produced during the training process for Model 3. The three lowest performing images past training image 700 have been highlighted in each case with their segmentations displayed in Figure 17. A total of 1130 images were included in training, but only 556 required corrective annotation (Table 1) due to inconsistencies in sponge presence and the success of Model 3.

| | RootPainter Prediction | Corrective Annotation | Metrics P | R | D | A | AE |
|---|---|---|---|---|---|---|---|
| A | 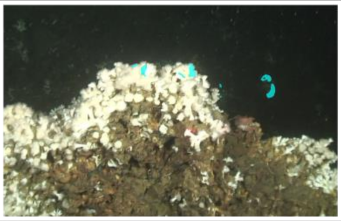 | 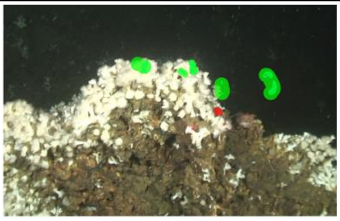 | 0.002 | 0.011 | 0.004 | 0.995 | 4.68 |
| B | 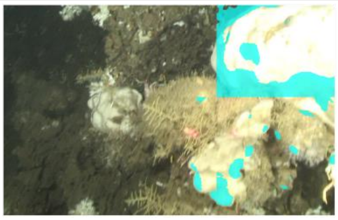 | 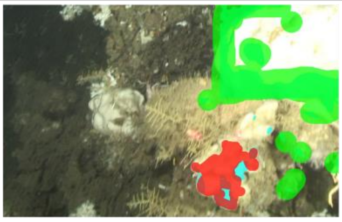 | 0.172 | 0.339 | 0.228 | 0.914 | NA |
| C | 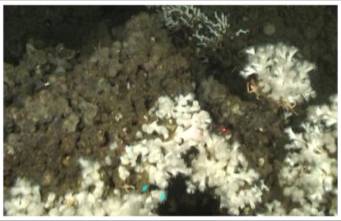 | 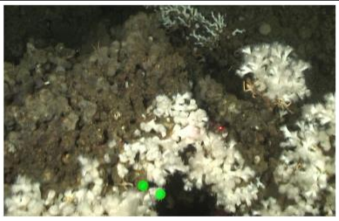 | 0.002 | 1 | 0.003 | 1 | 1.27 |
| D | 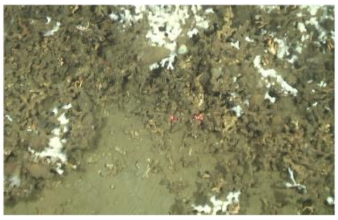 | 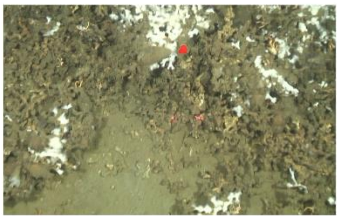 | 1 | 0.0528 | 0.100 | 0.999 | -1.98 |
| E | 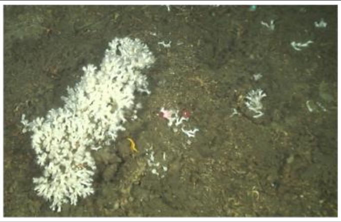 | 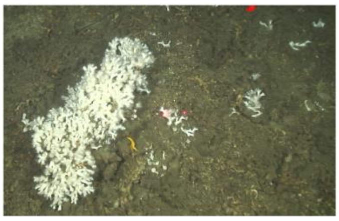 | 1 | 0.110 | 0.198 | 0.999 | -2.64 |
| F | 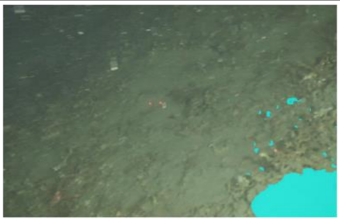 | 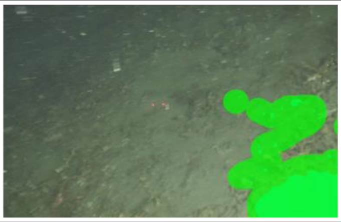 | NA | NA | NA | 0.951 | 813 |
| G | 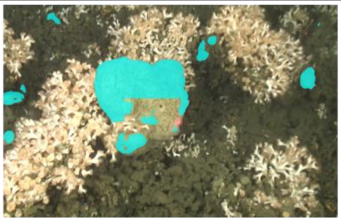 | 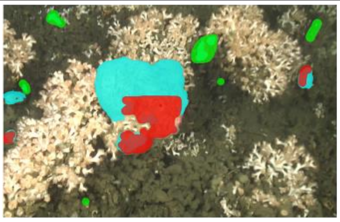 | 0.940 | 0.745 | 0.831 | 0.972 | NA |

SI Figure 6: RootPainter segmentations (light blue) and user corrections (with false positives annotated in green and false negatives annotated in red) relating to poor metrics from Figure 15. Where P = precision, R = recall, D = dice score, A = accuracy and AE = area error / $cm^2$. Where NA is present for area errors, Model 4 was not able to accurately segment the lasers for this image or only one laser was visible. Where NA is present for other metrics, the true/false positives/negatives required for their calculation are not present in corrections.

### Manual Validation Examples for Model 1

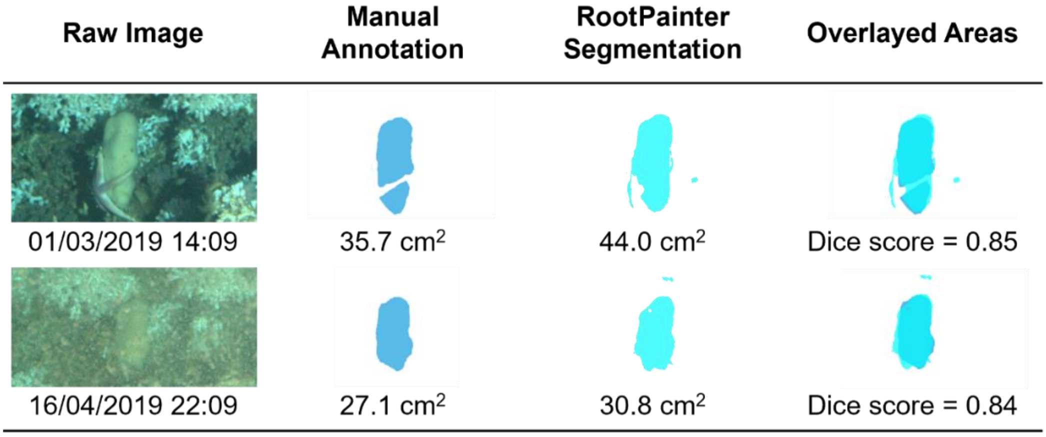


SI Figure 7: Images with ‘poor’ agreement between manual and RootPainter annotations.
